# Coexistence of self-compatible and self-incompatible alleles in a plant metapopulation via spatial rock-paper-scissors dynamics

**DOI:** 10.1101/2024.01.31.578129

**Authors:** Sachit Daniel, Mukund Thattai

## Abstract

Around 40% of flowering plants exhibit a preference for self-incompatibility systems (SI) on a macro-evolutionary scale. The dynamics of SI and self-compatible systems (SC) alleles can lead to the fixation of one genotype or coexistence, depending on factors like inbreeding depression and initial fractions. For the first time, we explore the dynamics of self-incompatible (SI) and self-compatible (SC) alleles in the presence of competitors, studying the relationship between population structure and SI frequency. Our numerical simulations consider non-instantaneous coupling between patches, focusing on a regime where the competitor’s fitness falls between inbred and outbred fitness. In this regime, SI, SC, and competitors exhibit repeating cycles of rock-paper-scissors dynamics both spatially and temporally. We find well-mixed environment leads to lower fractional population density of the focal species. Additionally in this regime, a paradoxical result emerges: in stress-free regions favorable to SI, the focal species may lose to the competitor.

## I. INTRODUCTION

Sexually reproducing organisms face competing selective pressures when choosing between selfing and outbreeding. Selfing alleles allow their pollen to fertilize their own eggs in addition to the eggs of any other individuals that will accept them. This increases the male reproductive success of selfers relative to outcrossers because outcrosser pollen can only fertilize the eggs of other individuals. However, in contrast to the selective pressure that favors selfing, the detrimental effects of reduced genetic diversity select for outcrossing. In the absence of any additional constraints, depending on the exact value of inbreeding depression, either self-compatible (SC) or self-incompatible (SI) strategies will be fixed in the population [1].

In plants, the presence of many flowers on the same individual creates an opportunity to self-fertilize (geitonogamy). To avoid self-fertilization, plants employ numerous physical and life-history mechanisms to reduce the amount of self-pollen deposited on the stigma.[2] In addition, numerous plant taxa have homomorphic molecular recognition mechanisms to identify and reject self-pollen at the cellular level. Around 39% of the flowering plant species have some mechanism of self incompatibility [3].

In this study, we focus on the gametophytic self-incompatibility (GSI) system. In GSI systems, a single, highly multiallelic locus (S locus) is used to mark, identify and then reject pollen from the same plant that produced it. They function by interrogating the haploid genotype of the pollen against both the alleles of the diploid parent plant (Fig 1a). If the S locus allele on the pollen has a functioning SI mechanism (I) and it matches with itself on the stigma, it is rejected. However unlike any of the self-incompatible alleles (I), pollen containing the self-compatible allele (C) at the S locus will not be rejected by the same plant and is free to self-fertilize. The possible fertilizations and the resultant genotypes of the GSI system are shown in Fig 1b. This dynamics is modelled by Eq 1. These types C alleles result in both the CC and IC individuals exhibiting an SC phenotype, while only individuals with an II genotype lead to SI.

**FIG. 1.**
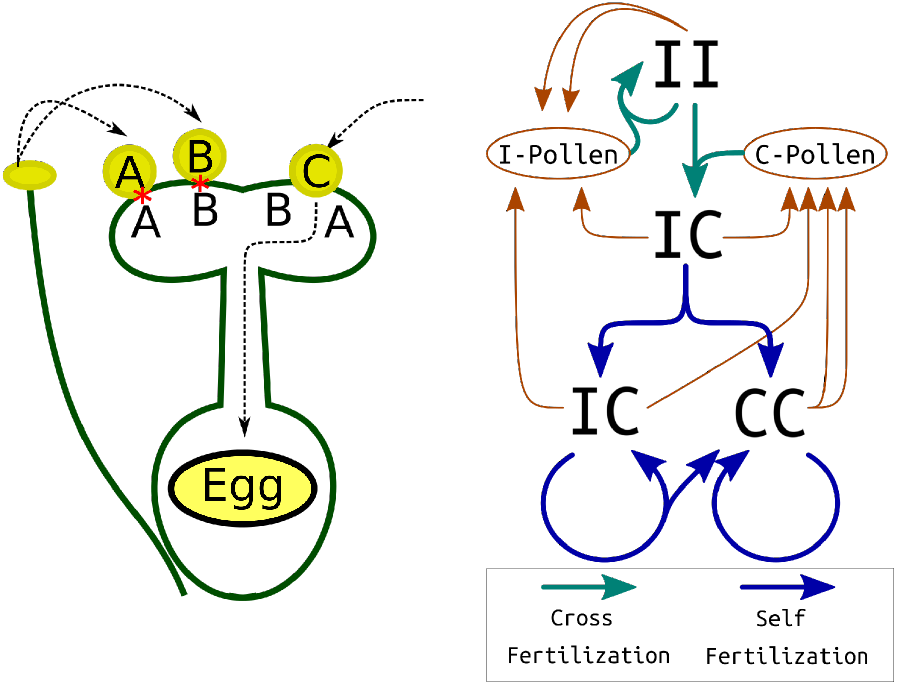
Gametophytic Self Incompatibility. **Fig 1a: Mechanism of Gametophytic Self Incompatibility (GSI)**. Haploid pollen expresses a single identity-determining motif interrogated by both alleles in the diploid maternal tissue. If the pollen matches either maternal determinant, fertilization is prevented. Pollen not matching either determinant is allowed to fertilize the egg. **Fig 1b: Model representation used in this study**. The self-compatible allele ‘C’ dominates over the self-incompatible allele ‘I’. Heterozygous individuals (IC genotype) are self-compatible, accepting their own ‘C’ pollen. All genotypes contribute to a pollen pool, and eggs of self-incompatible individuals (II geno-type) are fertilized with pollen of either genotype based on relative ratios in the total pollen pool.

SI is achieved through complex molecular cascades. Loss of function of certain genes in this cascade results in the formation of an SC allele [4]. In general, once most types of loss-of-function mutations are fixed, the restoration of function is difficult. While certain specific kinds of SI-SC mutations may be more easily reversible [5], analysis of the Solanaceae family has revealed that in practice, the transition from SI to SC is irreversible [6]. The repeated, irreversible transition from SI to SC by loss of function mutations and a subsequent selective sweep in the species should predict that the number of SC species must keep increasing relative to the SI species. However, the currently observed abundance of SI across many taxa is maintained by species level selection over a macroevolutionary timescale: SC species go extinct at a higher rate compared to SI species and SI species have a greater adaptive radiation into daughter species[7].

In this study, we explore the effect of a similar phenomenon, but at an ecological time scale. We explore the interaction between interspecific competition and the dynamics of SI-SC alleles. Natural habitats are tangled banks of multiple species with overlapping niches. Diversity is maintained by a number of *equalizing factors* (*sensu* Chesson, 2000 [8]) that bring the effective fitnesses of different species in an environment close to each other. Given that the effects of inbreeding depression is modulated by changes in the environment and competition [9][10], it should be expected that in some habitats, competitor species that are otherwise barely excluded from the niche of the focal species will have fitnesses higher than that of inbred individuals but still lower than that of an outcrossed individuals of the focal species. This study aims to examine the allele dynamics in such habitats where the fitness of a competitor species is bracketed by that of the outcrossed and selfed individuals of the focal species. We call this regime the rock-paper-scissors regime.

Previous work such as [11], [12] has shown that SI and SC alleles can coexist under certain circumstances. However they predict that coexistence will break down in the presence of very high selfing rates, low inbreeding depression and no sheltered load. This study takes these conditions where SI and SC are predicted to not coexist and see how it will interact with interspecific competition. Additionally, this study explicitly models the effects of limited dispersal and population structure to better mimic nature where ecological and evolutionary dynamics happens in the context of diverse interspecies interactions and across large geographical distances between the subpopulations of the global metapopulation of a species.

## II. THEORY AND METHODS

This study uses an individual-based simulation to analyze the problem of the coexistence of SI and SC alleles in a GSI system. We divide the environment into cells that can be occupied by either a single individual of the competitor species or by one of the genotypes of the focal species (see Table 1 for abbreviations and conventions). For the results presented in this paper, unless specified otherwise, the cells are arranged in a ring to model the perimeter of a contiguous environment. The simulation advances in discrete timesteps that correspond to generations.

**TABLE 1.** Terms, Abbreviations and Symbols.

| Focal species | The species in which we are examining the SI-SC dynamics |
| --- | --- |
| Competitor | A species that can exploit the niche of the focal species, but has lower fitness in that niche |
| GSI | Gametophytic Self Incompatibility |
| SI | Self Incompatible |
| SC | Self Compatible |
| I | Any of the SI alleles at the GSI locus |
| C | Any SC allele at the GSI locus |
| II,IC,CC | The three genotypes of the focal species, or their respective frequencies depending on context. II is shorthand for any heterozygote with genotype $I_x I_y$ where $I_x, I_y$ are each I alleles. |
| $F_x$ | The relative fitness of a genotype or species $x$, normalized to the fitness of the outbred focal species. |
| $\delta$ | Inbreeding depression. $F_{\text{Inbred}} = 1 - \delta$ |
| $p_I$ | The fraction of pollen that has an I allele |

In each generation, pollen containing I or C alleles is created from every individual of the focal species. This pollen is dispersed to the neighbouring cells within a certain radius proportional to a dispersal kernel. Specifically, we choose the kernel to be an exponential power distribution (a.k.a. the generalized normal distribution) with *β* = 0.4. This choice is based on the results of [13] who found that the heavy tailed generalized normal with a *β <* 1 best fits the observed dispersal in nature.

The SI plants of the focal species use this pollen to fertilize their eggs and create seeds. The inability of the SI plants to fertilize themselves is modeled by creating a notch at the center of the dispersal kernel: since the pollen from a plant can’t fertilize its own eggs, the situation is equivalent to the pollen never diffusing onto the source plant itself. The pool of I and C pollen that has landed on every cell is now used to fertilize the eggs of the focal species in the cell. SI individuals will fertilize their eggs with either C or I pollen in proportion to the quantity of the two types of pollen that have landed on the plant. The total amount of seeds that are produced by an SI plant is also a function of the total amount of pollen that has landed on the cell to simulate the effects of pollen limitation on seed set. For the results shown here, the an SI plant is assumed to have enough pollen to fertilize 99% of its eggs if it is within one *σ* of the pollen dispersal kernel of another conspecific. The SC plants are assumed to completely fertilize their own eggs: i.e prior selfing is modelled here instead of competing selfing or delayed selfing.

Note that while pollen discounting (reduction in the ability of selfers to disperse their pollen due to selfing adaptations) can be an important driver in the evolution of self incompatibility [14], in this study, we assume that there were no *marginal* pollen discounting effects in the SC plants because the C alleles that are under examination act via molecular recognition of previously deposited pollen. Non-homomorphic traits, such as variations in flower morphology, may simultaneously affect the rate of selfing and the level at which selfing reduces the spread of pollen to other plants. For example, they may affect the amount by which the pollen deposited on a pollinator is removed by the stigma of the same flower. However, the mutations in the molecular recognition pathway that act solely post-pollination have no marginal effects on the pollen dispersal.

Each individual of the focal species is marked as inbred or outbred based on whether they are a product of selfing or outcrossing. Inbred individuals could suffer a penalty in the amount of seeds and pollen they produce or in the level of recruitment to the next generation. Unlike in previous work [11], we choose to model inbreeding depression as a reduction in the overall growth rate instead of modeling it as a process that is driven primarily by recessive lethals affecting germination of the seeds [15].

A simplifying assumption is made that the level of inbreeding depression will not compound over generations: the reduction in fitness of an inbred plant compared to an outbred plant is assumed to be the same irrespective of whether the plant is derived from one generation or from multiple generations of selfing. Additionally, the depression is assumed to be resistant to purging (i.e. successive generations of inbreeding cannot completely eliminate the deleterious alleles responsible for depression) and any depression that has accumulated due to inbreeding can be completely restored after outcrossing (i.e. outcrossed offspring of inbred parents have the same fitness as other outcrossed individuals).

Biologically, these approximations have empirical justification [16, 17] and will be caused by the depression being driven by numerous loci, with each locus having a small effect that are under selection for heterozygosity. This could be because of either natural selection directly favouring diversity of alleles at that locus (overdominance) or because there are weak deleterious mutations that are linked in repulsion (pseudo overdominance)[17–19]. Even in the absence of pseudo overdominance, if inbreeding depression is caused by mutations of small effect it will be resistant to purging. Previous work has shown that when weak effect mutations are explicitly simulated, the difference in behaviour due to approximating inbreeding depression as fixed is negligible.[20] Empirical work has shown that the effects of inbreeding does not show up as large effect mutations that limit the viability or germination of the seeds of plants, but instead acts later in life.[15] Recent theoretical work has also suggested that the presence of multiple types of mutations that act at different life stages will interact with each other to interfere with the purging of late acting small effect mutations [21, 22].

The offspring produced by selfing will have reduced fitness *F*_Inbred_ which is the same as 1 − *δ*, where *δ* is the inbreeding depression. This term captures all the effects starting from the fertilization to the next gametogenesis, i.e., the product of reduction in seed-set, survival, and recruitment, or it can be used to capture the effects of reduction in post recruitment biomass and flowering. We use this form of representing the depression instead of the conventionally used *δ* because it makes it easier to compare its interaction with the fitness of the competitor species (*F*_Competitor_) since that is the focus of this study.

Next, the seeds produced by each plant are dispersed to their neighboring cells within a certain range via a seed dispersal kernel which is also chosen to be an exponential power distribution with *β* = 0.4 [13]. For the results shown below, the seed dispersal radius was set to be one third that of the pollen dispersal radius, though the results are qualitatively robust to variations in this ratio. The dispersion radius determines how well-mixed the environment is; specifically, the smaller the dispersal radius relative to the total environment, the less well mixed the environment is.

In every generation, there is a probability that a small number of seeds of any type is added to the environment at a random location to represent the effects of immigration from a different environment that is not contiguous with this one and has only a weak level of coupling. Since we do not explicitly model the presence of S alleles and the genetic architecture of inbreeding depression, we have taken care to use this pulsed immigration (resembling variations in faraway populations) instead of immigration of individual seeds. This is so that any results that we observe will not be confounded by colonization happening with just two SI seeds which will produced a bottle necked population that falls outside the approximations of the model.

After each generation, the plants in each cell are replaced with a new individual randomly recruited from the pool of seeds that have landed in that cell with a probability proportional to the density of that type of seed.

## III. RESULTS

The description of the rock-paper-scissors dynamics for a single cycle is presented in section III A. Then, we discuss how this cycles reinitiates to repeat itself in section III B. Finally, we describe how the repeated cycles develop temporal and spatial structure in section III C.

### A. Dynamics in a single Rock-Paper-Scissors cycle

In our study we focus on the regime where the competitor’s fitness is bracketed by the fitness of the outbred and the inbred individuals of the focal species: i.e., *F*_Outbred_ *> F*Competitor *> F*Inbred *> F*Threshold. In this regime the dynamics goes as follows: because *F*_Outbred_ *> F*_Competitor_, a purely SI (and hence outbreeding) population of the focal species can invade and replace the competitor. Additionally, since *F*_Inbred_ *> F*_Threshold_, any introduction of a C allele to the population of the focal species will result in the I allele being driven extinct and the focal species becoming purely SC. However, because *F*_Competitor_ *> F*_Inbred_, a population of pure SC focal species will be vulnerable to invasion by the competitor. This results in the pure states of either SI(II), SC(CC) or the competitor being unstable because an invasion of a small number of individuals with a superior strategy can causes the population to move to a different state. We call this regime of cyclic dominance the “rock-paper-scissors” regime. A classical rock-paper-scissors system is characterised by a cyclic dominance between the different players where all three players have positive and negative frequency dependant effects on each other. In this case, the population will undergo sustained oscillations between the different species [23]. However, in our system there are no asymmetric frequency dependent relationships between the competitor and either genotype of the focal species. Due to this, any mixed population of all three types will eventually converge to a pure competitor state. Unlike the classic system of rock-paper-scissors, the dynamics needs to be reinitiated and we discuss this mechanism in section III B.

The dynamic in a single cycle of rock-paper-scissors happens as follows: the effective growth rate of SI individuals is reduced in proportion to the allele frequency of C, while the growth rate of SC individuals is enhanced by the same amount. In each generation, an increasing fraction of the SI is converted to outbred SC individuals with the same fitness as the SI. In a generation after SI has be driven to extinction, all SC will be inbred. While the competitor cannot outgrow the outbred SI and outbred SC of the focal species, it will outcompete inbred SC due to the difference in their fitness. Once sufficient inbreeding depression sets in, the patch is replaced by a competitor that was held in check for its mild disadvantage compared to the outbred individuals of the focal species. Eventually, in a finite population, SC (and hence the focal species) will be driven extinct. We see this dynamics in our numerical simulation outlined in section II.

### B. Re-ignition of the Rock-Paper-Scissors cycle

When the patch is completely occupied by the competitor, it can eventually be invaded by SI individuals. Since the fitness of SI is greater than that of the competitor, SI will replace the competitor gradually, and reset the cycle described above. Note that only the pure competitor state is vulnerable to being invaded by SI and is a necessary condition for re-ignition. In a patch which is in the rock-paper-scissors fitness regime, if there are any remaining SC individuals of the focal species e.g., due to insufficient time passing for the cycle to complete, or due to sufficient external migration that continuously repopulates SC, the patch will be resistant to the invasion of SI. Since the inbred focal species will not be able to grow faster than the competitor, the population of the focal species will remain suppressed and the cycle will not re-initiate. However, note that if external migration is low enough that stochastic variations allow the local population to be extirpated, a small introduction of SI will lead to the re-initiation of the cycle. When the radius of dispersal is small relative to the total size of the contiguous environment, the metapopulation behaves like a set of patches that are only weakly coupled. If the dispersal radius is small, then stochastic fluctuations in the distribution of individuals will cause the recolonization of SI individuals to occur in a patchy manner, instead of being contiguous with the rest of the population. This leads to coexistence of SI and SC alleles across the metapopulation.

In Fig 2a, we present the waterfall plot of a slice of the population over time. The repeated cycles of colonization and extirpation that occur in this dynamics can be seen this Fig as vertical stripes (i.e, a choice of patches), undergoing the above sequence of replacements. Fig 2c shows the average temporal cross-correlation of relative density of various genotypes in the metapopulation as a function of the population of outbred SC individuals. Moreover, due to the stochastic nature of the recolonization of new patches, we expect no long range (i.e, range much greater than dispersal radius) spatial pattern to the distribution of patches over time. This is because different parts of the environment will be undergoing different phases of the cycle independently. This can be seen in Fig 2b.

**FIG. 2.**
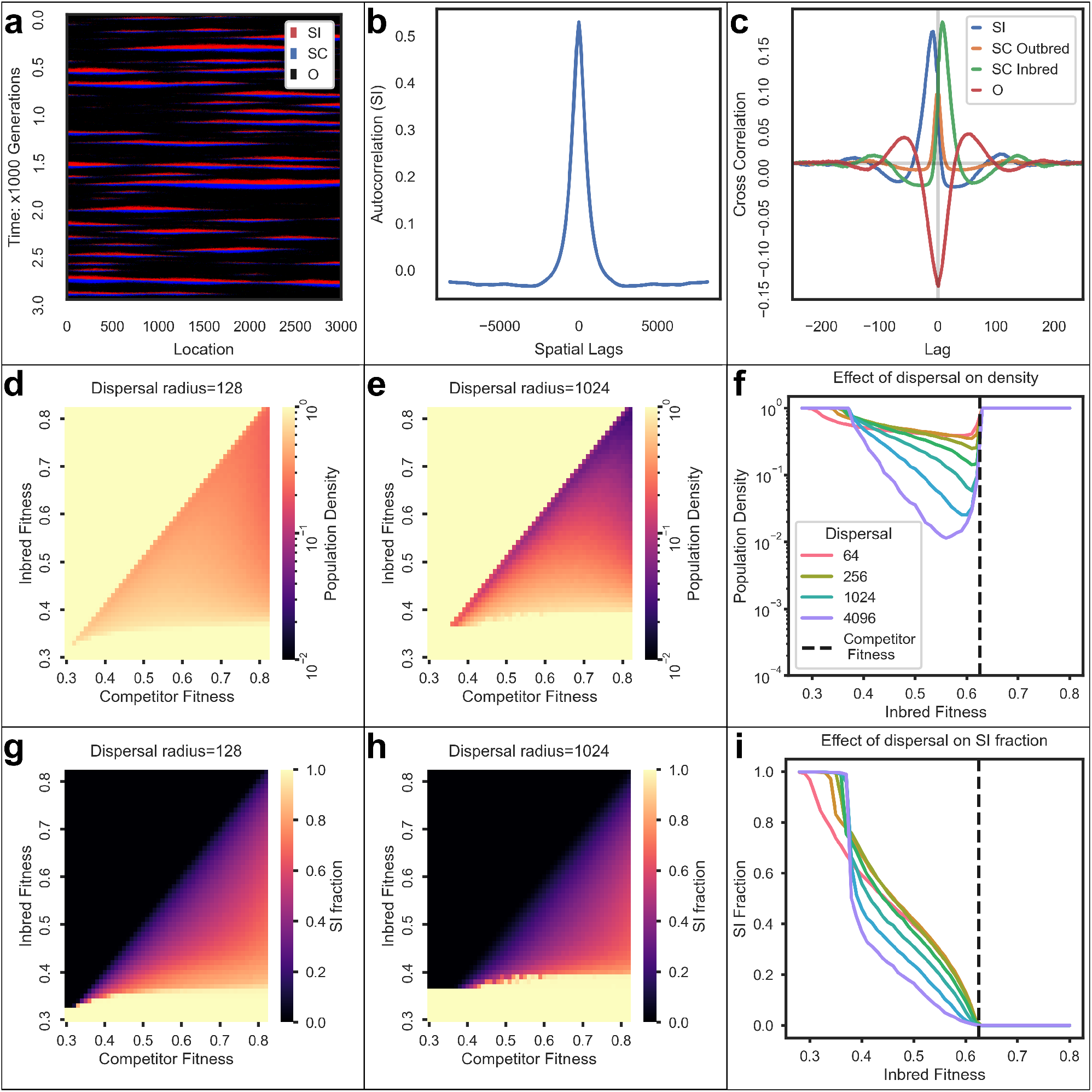
Row 1: Coexistence at a metapopulation level driven by out-of-phase succession dynamics at a sub-population level. **Panel a**: The waterfall-plot illustrating the population dynamics for dispersal radius of 512. The competitor (black) is replaced by SI (red), succeeded by SC (blue), and then the competitor returns. **Panel b**: Short-range correlation in the spatial auto-correlation of the SI population, without long-range periodicity. **Panel c**: Temporal cross-correlation of relative population density, highlighting a characteristic local succession dynamics without long-range temporal correlations. Patches cleared of competitors are invaded by SI, succeeded by SC, and eventually replaced by the competitor. **Row 2: High dispersal and relaxed selection can reduce the population density of a species**. Below *F*_threshold_, SI alleles dominate, excluding competitors. In the range *F*_threshold_ *≤F*_inbred_ *≤F*_competitor_, SI and SC alleles coexist through *rock-paper-scissors* dynamics, reducing the focal species density. Above *F*_inbred_ *≥ F*_competitor_, the focal species excludes competitors, comprising SC-only individuals. Increased dispersal approximates well-mixed system dynamics without coexistence. **Panels d and e**: Population density of focal species for two fixed dispersal radii **Panel f** : Average focal species density against inbred fitness. **Row 3: Effect of dispersal radius on allele coexistence**. Limited dispersal radius and *F*_outbred_ *≥ F*_competitor_ *≥ F*_inbred_ *≥ F*_threshold_ lead to allele coexistence; otherwise, either SI or SC is fixed. **Panels g and h**: Fraction of the focal species for two fixed dispersal radii. **Panel i**: Fraction of the focal species at various dispersal levels with *F*competitor fixed at 0.825.

### C. The temporal and spatial structure of the Rock-Paper-Scissors cycle

In Fig 2f, we show the fractional population of focal species i.e., 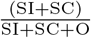 as a function of inbred fitness for various dispersal radii holding the fitness of the competitor fixed. Note in these plots the quantities are time and space averaged and *F*_outbred_ is set to 1. In our simulation, we see that as the dispersal radius increases, the total fraction of focal species decreases. The decrease in the fraction of the focal species with an increase in the dispersal radius is caused by two processes:

a. Within a contiguous environment, if the radius of dispersal is large for a given level of migration, it will be harder to find a patch that is entirely cleared of the focal species. In contrast, with a smaller dispersal radius, it will be easier to find empty patches that can re-initiate the cycle. This allows the SI alleles to establish a population in areas that have been cleared of the focal species without immediately being outcompeted by the SC alleles. Despite larger dispersal radii resulting in a greater number of re-seedings of SI individuals, the number of productive reseedings leading to a successful invasion is lower. This is attributed to the fact that a proportionately larger patch must be cleared before establishing a zone free of SC pollen, causing the cycle to occur at a slower pace. This will result in the average population of the focal species being reduced due to a delay in the colonisation-expansion phase.
b. The second process acts after a successful colonization because a patch containing the focal species can only grow until it couples with another patch containing SC individuals, after which the population flows toward a competitor-only state. The entry of an external SC allele into a patch is facilitated with a larger dispersal radius, causing the collapse of the focal species to begin sooner.

Next, we show the interaction between *F*_competitor_ and *F*_inbred_ on the fractional population of the focal species in Fig 2d-e. We choose to depict two representative radii corresponding to 128 and 1024 cells (note that the total size of the environment is 16,384 cells) but confirm that the qualitative pattern is robust to the choice of radius. Notice that as the dispersal radius increases the population density goes down. The two simple cases are as follows: above the diagonal (*F*_competitor_ *< F*_inbred_) in 2d-e the competitor is excluded, and when *F*_inbred_ *< F*_threshold_, we have a focal species-only state because SI dominates. In the region where *F*_competitor_ *> F*_inbred_ *> F*_threshold_ (i.e., the rock-paper-scissors regime), we see that as *F*_competitor_ and *F*_inbred_ increase, the population of the focal species declines. This is because higher *F*_inbred_ leads to SI being replaced by SC sooner followed by the invasion of the competitor. As *F*_competitor_ increases, the SI re-invasion becomes harder. We also see this playing out when we look at a horizontal strip (i.e., holding *F*_inbred_ constant). The population of the focal species is greatest when there is a competitor that is fit enough to purge the inbred individuals, but not so competitive that the SI cannot re-establish itself.

In third row of Fig 2, we examine the the factors influencing the SI fraction of the focal species. In Fig 2g-h, the relative fraction of SI monotonically decreases with *F*_inbred_ and increases with *F*_competitor_. This is because of the reduction in the time spent where the SC individuals dominate the population. Comparing panels g and h, we see that SI faction increases with increasing dispersal radius. In Fig 2i, we investigate the variation in the relative fraction of SI with changes in *F*_inbred_ and dispersal radius, while keeping *F*_competitor_ constant. As expected from the process discussed in the previous section, an increase in the dispersal radius leads to a decrease in the relative fraction of SI. As the dispersal radius increases, the population starts to behave like a well-mixed system wherein the coexistence of the two alleles is reduced.

The overall interaction of the effects of dispersal rates, inbreeding depression, and competitor fitness are summarized in Fig 4. Lastly, it’s important to highlight that at low dispersal radii, the threshold for SC invasion is decreased. Further details on this is elaborated in the supplementary material.

### D. In the presence of environmental cline in fitness

High migration rates can suppress the population of the focal species as discussed in the previous section. One source of a high migration rate of SC into a rock-paper-scissors environment will be where there is a cline in the environment leading to the rock-paper-scissors regime bordering an area where the focal species can grow stably. We examine a case where there is a cline in the fitness and depict the results in Fig 3. At the point in an environmental cline where the fitness transitions from a regime of *F*Threshold *> F*Inbred to *F*Inbred *> F*Threshold, population will transition from being dominated by SI to SC. However, here SC alleles will be present primarily as outbred individuals formed by the fertilization of SI individuals with SC pollen. A short distance away from this transition zone there will be a patch in which the population of the focal species is suppressed by the competitor. This happens because outbred SC individuals that migrate from near the transition region with fitness similar to SI will prevents the SC allele from being fully purged. The presence of the SC individuals will then prevents an out-breeding SI population from taking hold. However, since inbred SC individual’s fitness is smaller than the competitor, they are eliminated by the competitor. This “wall” of SC individuals will spatially curtail the focal species’s ability to expand into a patch where it is nominally more fit than its competitors. Comparing the two panels in Fig 3, it can be seen that the rock-paper-scissors interaction causes the focal species to be suppressed by the competitor.

**FIG. 3.**
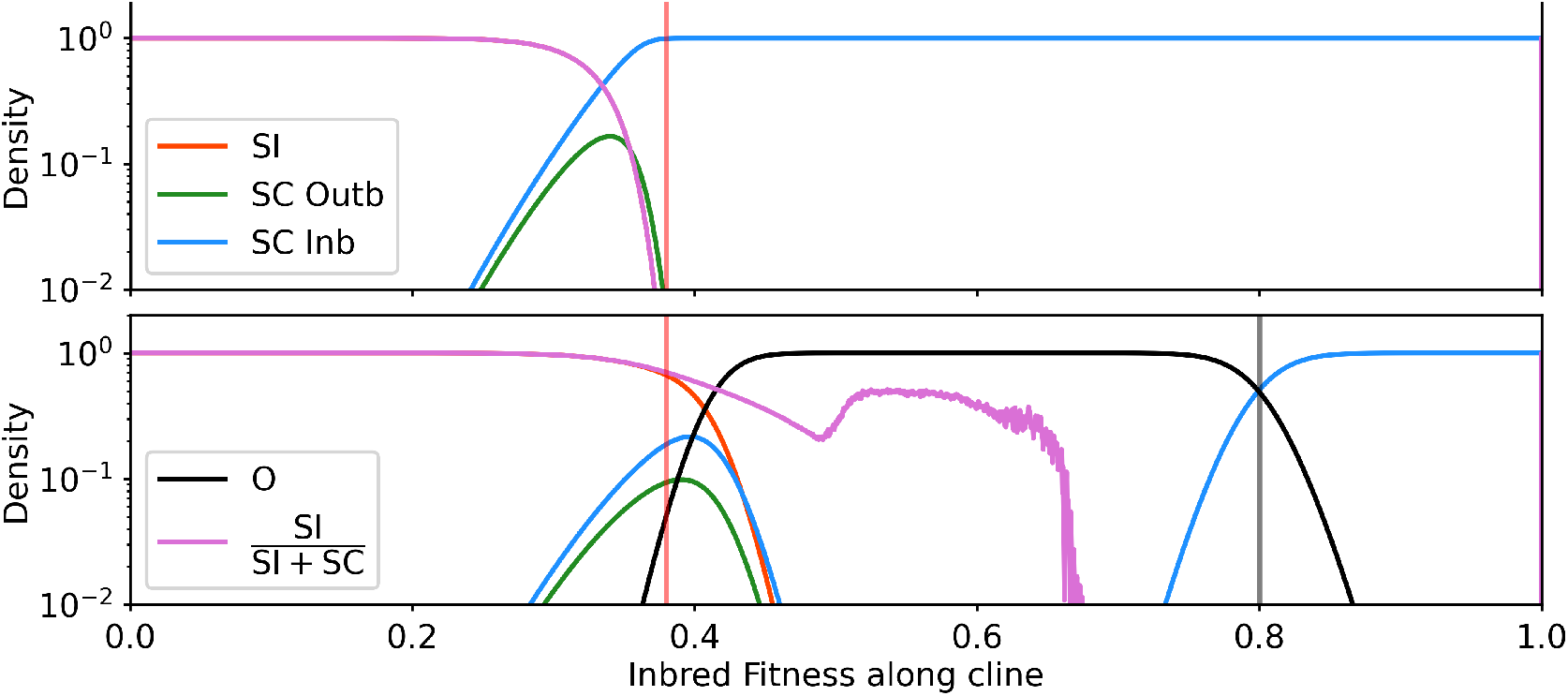
SC alleles generate a barrier that prevents range expansion: This figure shows the population composition in an environment with a fitness gradient. Going from the left edge of the environment to the right edge the fitness of the inbred individuals increases from below the threshold to above that of the competitor. At the border between a permissive and unfavourable environment, SC alleles exist primarily as outbred individuals in. Further into the permissive environment where the inbred individuals are fitter than the threshold, the population is completely dominated by SC individuals. However, the competitor species can completely eliminate the focal species. The constant influx of SC pollen from the barrier prevents SI individuals from recolonizing the niche and displacing the competitor. The focal species dominates and gets reestablished only when the fitness exceeds that of the competitor. The two panels panels show the effect the absence (upper panel) or presence (lower panel) of a competitor can have in reducing the range of the species and increasing the apparent frequency of self incompatibility.

**FIG. 4.**
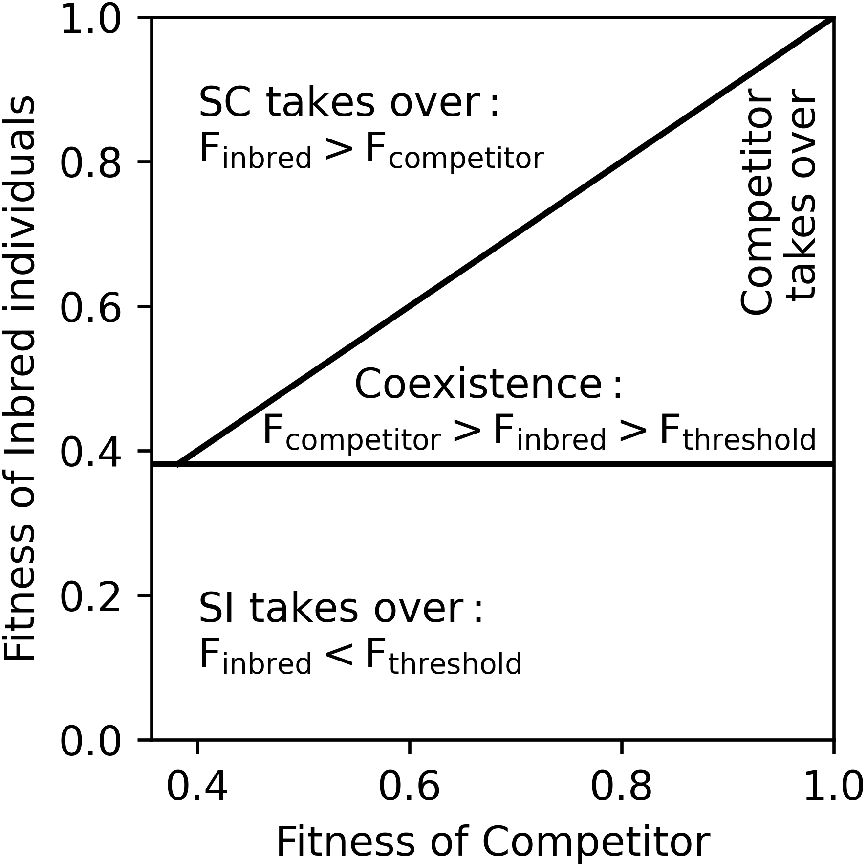
A simplified representation of the dynamics of a GSI system in the presence of competitors and limited dispersal. When the condition *F*_outbred_ *≥ F*_competitor_ *≥ F*_inbred_ *≥ F*_threshold_ is satisfied, there is coexistence of the two alleles. Outside of this range, either SI or SC is fixed. Within the zone of coexistence, as *F*_inbred_ approaches *F*_outbred_, the population density of the focal species drops and the competitor starts to dominate.

## IV. DISCUSSION

Empirical data has shown that there is a great deal of intra-species variation in the level of selfing across space and time [24, 25]. While most theories predict that SI and SC strategies will coexist only for a narrow range of parameters [11, 12], we provide an additional mechanism that might contribute to the observed pattern of mixed mating strategies. Even in the presence of a temporally and spatially homogenous selection regime, the cyclic dominance between SI, SC and competitors can create large variations in both the population size and selfing rates when measurements are carried out at a scale much larger than the dispersal radius of the plants.

On the surface, the metapopulation results of our model go against the intuitions one would get from bakers law: i.e., lower dispersal rates should be expected to lead to greater mate limitation, which will in turn lead to SI increasing instead of decreasing with well mixed environments. The reason for this deviation from intuition is that, in our model we assume that immigration happens in pulses of immigrants with a low, uniform probability of a point in the perimeter of the simulated island getting the pulse of immigrants. This will biologically correspond to the stochasticity in the immigration being driven by dynamics in environments far away from the focal environment. If a much lower rate of immigration is modelled by assuming that the seeds land as individuals, for a low enough migration probability, the system starts to replicate the predictions of bakers law, where the lower dispersal rates lead to greater SC instead of SI due to mate limitation.

Recent work has revealed that there is no general trend of selfing rate across populations of species when going from the central to the peripheral zones[26]. However, they did reveal that there was significant heterogeneity between species on this metric.

In theory, numerous processes have been hypothesised to increase selfing towards the margins of a species range. Selfing can expand a species range through a variety of ecological and bio-geographic factors. Beyond the optimal habitat of the species, Allee effects (through absolute mate limitation or pollinator limitation) can sharply curtail the ability of SI individuals to establish themselves. SC allows species to maintain a foothold in or propagate a colonization wave into new habitats. In addition to these effects, it has been proposed that gene flow and outcrossing with a core population can reduce the ability of adaptive gene combination from getting fixed in peripheral zones. Additionally, certain kinds of stresses can also reduce the in-breeding depression by capping the growth of plants indiscriminately. These mechanisms all predict that the margins of the species range *cause* an enrichment of selfing.

In contrast to this effect, we find in this study that selfing can cause the limits of the species range by greatly curtailing the habitable range of the species. In unfavourable environments that reduce the fitness of the species, the species can be driven extinct or its population be greatly suppressed even if its fitness isn’t reduced below any competitor’s. The key factor lies in stressors imposing just enough constraint on the growth, thereby reducing the disparity between SI and SC fitness. This reduction needs to be just sufficient for SC to surpass the threshold *F*_threshold_. Our model makes another counter intuitive prediction: In highly permissive habitats, with no stressors to challenge inbred plants, in-breeding depression can also be reduced. This will also lead to the rock-paper-scissors dynamics suppressing the population of the species. Taken together this model predicts that there may exist a Goldilocks level of stress for a species to thrive in a tangled bank. This model predicts that relatively small disturbance to the environment (e.g., climate change, habitat fragmentation) can have a disproportionate impact on the population. These human induced perturbations need not necessarily directly harm the species; even alteration that nominally increase the growth rate e.g., relieving nutrient limitation can cause the population to collapse.

## V. SUPPLEMENTARY MATERIALS

In this section, we extend the study beyond the previously explored rock-paper-scissors regime. Unlike the main paper where we concentrate on the region where the fitness of inbred individuals (*F*_inbred_) exceeds the threshold (*F*_threshold_ ∼ 0.38), here we relax this criterion.

First, let’s examine the model without the presence of the competitor in a region whose size is comparable to the dispersal radius. This small region of the environment can be treated as approximately well mixed. In these patches, our system’s dynamics can be analytically described using the equation 1 which is a special case of the model in [11].

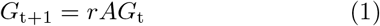

where,

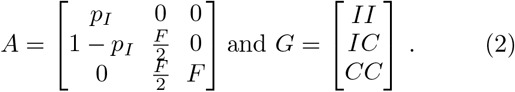

Here, *r* is the population growth rate of a population composed entirely of II and *r* × *F* is the growth rate of a population composed entirely of CC. Among the three characteristic equations of *A*, two are trivial. The third solution, however can be utilized to establish the *F*_threshold_ which turns out to depend on the initial conditions of the population as follows:

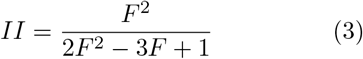

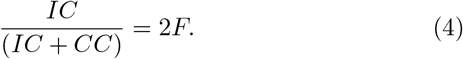

We present the set of curves corresponding to different initial conditions in Fig 5.

**FIG. 5.**
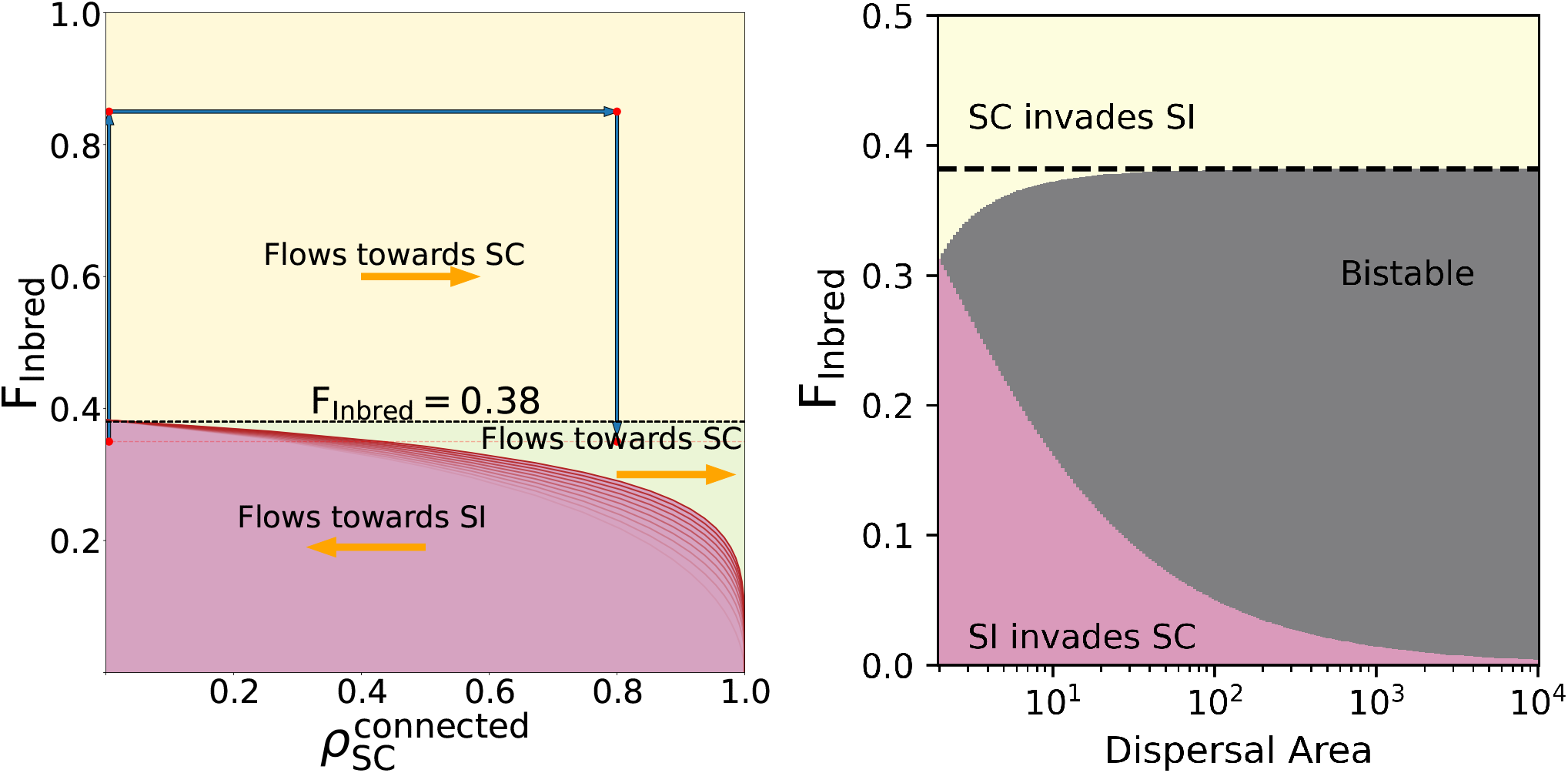
Left panel: The phase-space diagram and hysteresis: This figure provides an overview of different regions in the *F*_inbred_ and 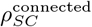 phase-space, detailing the dynamics within them. The red curves correspond to different initial values of SC homozygosity. The system flows toward an SI-only population below these curves and toward an SC-only population above them. The yellow region, emphasizing repeated rock-paper-scissors dynamics, is the primary focus of this paper. The red dots along with the blue arrows illustrate the concept of hysteresis in this phase space, where the system, with the same *F*_inbred_ value, transitions to either an SI-only or SC-only state depending on its past condition. **Right panel: Limited dispersal reduces bistability in GSI**: GSI systems show a region of bistability where either SI or SC are evolutionarily stable. When dispersal is reduced, the increase in the local concentration of small perturbations causes the system to lose it’s zone of bistability.

In the phase-space defined by *F*_inbred_ and 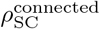 (the local concentration of SC in a well connected patch), the curve provides the value of *F*_threshold_ as a function of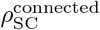. The system’s dynamics transition around curves and converge towards one of two fixed states. These states are characterized by either the entire population in the connected patches being exclusively composed of SCs or solely composed of SIs. More specifically, if the system lies above this curve, it will ultimately reach a fixed point exclusively composed of SCs; conversely, if it lies below this curve, the system will eventually reach a fixed point exclusively composed of SIs. In this case, we see that there is no coexistence of SC and SI strategies.

Next, we introduce the competitor species to this system. The interaction dynamics between SC and SI individuals remain independent of the competitor’s presence. This distinction from a classical rock-paper-scissors setting is highlighted in III. The competitor’s impact on the dynamics lies in determining whether, once the fixed point mentioned earlier is reached, the competitor can either replace or fail to replace the focal species. In the presence of a competitor whose fitness falls between that of the inbred and outbred focal species, reaching the fixed point occupied by SC individuals results in the replacement of the focal species by the competitor.

Now, the phase space can be divided into three regions, as indicated in the left panel of Fig 5. The area below the curve (highlighted in pink) corresponds to the fixation of SI. While SC can invade and disrupt this state, a sufficiently large number of SC is needed to cross the curve and flow into the other fixed state. The system therefore stabilizes at the SI fixed point and is robust against small perturbations (low levels of SC introduction). This state is also resistant to competitors.

We can partition the region above the curve, where the fixed point in the absence of a competitor is a population of SC, into two segments: one where *F*_inbred_ *>* 0.38 (highlighted in yellow) and another where *F*_inbred_ *<* 0.38 (highlighted in green).

In the *F*_inbred_ *<* 0.38 region (highlighted in green), while a purely SC state is resistant to the invasion of SI, the competitor can still replace the inbred SC. Once the competitor entirely replaces SC, it in turn becomes susceptible to invasion by SI from various patches in the environment. SI takes over, resulting in a state of an exclusively SI population. The system has now entered the state in the pink region where it is stable to the introduction of competitors or small numbers of SCs. Thus, while it is possible for the system to get stuck in an SC-only state even with high inbreeding depression (green), the presence of competitors allows a patch of focal species to “reset” to the SI state in all cases with *F*_inbred_ *<* 0.38. This demonstrates a behaviour at the ecological scale that mimics that which is seen in a macroevolutionary timescale.

Contrasting this, the last region highlighted in yellow, where the *F*_inbred_ *>* 0.38 is the region where we have the repeated cycles of rock-paper-scissors. This region is the focus of our paper and has been explored in length using numerical simulations of the system in section III.

The concentration dependant effects that have been described above can be extended to analyse the effect of limited dispersal on GSI systems. When dispersal is limited, even a small number of individuals of any type will have the effect of a much higher concentration locally. In the right panel of Fig 5, we illustrate the system’s behaviors with respect to the dispersal area measured in the number of individuals and *F*_inbred_. When *F*_inbred_ *>* 0.38 (the green region), SI is unstable to even a small introduction of SC and the system dynamics enables SC invasion and fixation. If a suitable competitor is introduced, this region will exhibit the rock-paper-scissors dynamics discussed in the paper.

In the limit of a well-mixed system, i.e., a large dispersal area, when *F*_inbred_ *<* 0.38 (the grey region), the focal species exhibits bistability. It can evolve into either an exclusively SI population or an exclusively SC population, and these states remain robust against small introductions of SI or SC. An effect of the bistability is that once SC is fixed in an island through any means, it will be very hard for SI to invade. However, if a suitable competitor is introduced, it will disrupt the SC stable state and allow SI to invade. Once SI has taken over, it will now be resistant to the invasion of SC. This effect can be seen in the very bottom of Fig 2g-h, and the very left of Fig 2i, where the population is entirely SI.

If the dispersal area is small, the region can be delineated into two zones. In the region where 0.31 *< F*_inbred_ *<* 0.38 (depicted in green), the region of bistability is reduced allowing small introductions of SC to invade an SI population. The value of 0.31 comes from the *F*_inbred_ needed to prevent SC from invading when it is present in an equal amount as SI in the population from Eq. 1. When the dispersal radius is small, small amounts of invading SC will experience a high local concentration irrespective of how large the entire environment is. At the “front” of an SC invasion, the SI and SC individuals will be present approximately equally. This extends the zone of rock-paper-scissors dynamics in the presence of a competitor and can be seen in the increased coexistence for low dispersal radii in Fig 2i. While it has been previously shown[20, 27] that limited dispersal will negatively affect the stability of SI through reduction in the diversity of S alleles and by effectively reducing the marginal inbreeding depression of SC due to the effects of bottlenecks, we show that there is an additional effect that will destabilise SI at low dispersal radii.

Finally, when dispersal is low and *F*_inbred_ *<* 0.31 (shown in pink), the reduced bistability allows SI to invade even if SC is already fixed resulting in a stable SI-only population even without the presence of a competitor. However, this result may not be biologically relevant because low dispersal rates will disproportionately penalise SI by its effects on the diversity of S alleles and in the rest of the genome. However, as shown in the rest of the paper, the presence of competitor reduces the burden on SI’s ability to directly replace an SC population.

## VI ACKNOWLEDGEMENTS

This work was supported by the Simons Foundation (287975 to MT). SD thanks Swetha Bhagwat for her invaluable help in turning what was just a pile of results into an actual paper. SD thanks Shweta and Terence for all the support they have given through this work.

## VII. AUTHOR CONTRIBUTIONS

MT conceived the study. SD developed the model and performed the analysis. SD and MT wrote the paper.

## References

[1] D. Charlesworth and B. Charlesworth. The evolution and breakdown of S-allele systems. Heredity, 43(1):41–55, 1979. ISSN 1365-2540. doi:10.1038/hdy.1979.58. URL https://www.nature.com/articles/hdy197958.

[2] Vernonica E. Franklin-Tong. Self-Incompatibility in Flowering Plants. Springer Berlin Heidelberg, Berlin, Heidelberg, 2008. ISBN 978-3-540-68485-5978-3-540-68486-2. doi:10.1007/978-3-540-68486-2. URL http://link.springer.com/10.1007/978-3-540-68486-2.

[3] Boris Igic, Russell Lande, and Joshua R. Kohn. Loss of self-incompatibility and its evolutionary consequences. International Journal of Plant Sciences, 169(1):93–104, 2008. ISSN 1058-5893. doi:10.1086/523362. URL https://www.jstor.org/stable/10.1086/523362. Publisher: The University of Chicago Press.

[4] Jianke Du, Chunfeng Ge, Tingting Li, Sanhong Wang, Zhihong Gao, Hidenori Sassa, and Yushan Qiao. Molecular characteristics of s-RNase alleles as the determinant of self-incompatibility in the style of fragaria viridis. Horticulture Research, 8(1):1–18, 2021. ISSN 2052-7276. doi:10.1038/s41438-021-00623-x. URL https://www.nature.com/articles/s41438-021-00623-x. xNumber: 1 Publisher: Nature Publishing Group.

[5] Sota Fujii, Ken-ichi Kubo, and Seiji Takayama. Non-self- and self-recognition models in plant self-incompatibility. Nature Plants, 2(9):1–9, 2016. ISSN 2055-0278. doi: 10.1038/nplants.2016.130. URL https://www.nature.com/articles/nplants2016130. xNumber: 9 Publisher: Nature Publishing Group.

[6] Boris Igic, Lynn Bohs, and Joshua R. Kohn. Ancient polymorphism reveals unidirectional breeding system shifts. Proceedings of the National Academy of Sciences, 103(5):1359–1363, 2006. doi:10.1073/pnas.0506283103. URL https://www.pnas.org/doi/full/10.1073/pnas.0506283103. Publisher: Proceedings of the National Academy of Sciences.

[7] Emma E. Goldberg, Joshua R. Kohn, Russell Lande, Kelly A. Robertson, Stephen A. Smith, and Boris ć. Species selection maintains self-incompatibility. Science, 330(6003):493–495, 2010. doi: 10.1126/science.1194513. URL https://www.science.org/doi/10.1126/science.1194513. Publisher: American Association for the Advancement of Science.

[8] Peter Chesson. Mechanisms of maintenance of species diversity. Annual Review of Ecology and Systematics, 31 (1):343–366, 2000. doi:10.1146/annurev.ecolsys.31.1.343. URL https://doi.org/10.1146/annurev.ecolsys.31.1.343.

[9] Cheptou, Imbert, Lepart, and Escarre. Effects of competition on lifetime estimates of inbreeding depression in the outcrossing plant crepis sancta (asteraceae). Journal of Evolutionary Biology, 13(3):522–531, 2000. ISSN 1420-9101. doi:10.1046/j.1420-9101.2000.00175.x. URL https://onlinelibrary.wiley.com/doi/abs/10.1046/j.1420-9101.2000.00175.x.

[10] Tobias M. Sandner, Diethart Matthies, and Donald M. Waller. Stresses affect inbreeding depression in complex ways: disentangling stress-specific genetic effects from effects of initial size in plants. Heredity, 127(4):347–356, 2021. ISSN 1365-2540. doi:10.1038/s41437-021-00454-5. URL https://www.nature.com/articles/s41437-021-00454-5. xNumber: 4 Publisher: Nature Publishing Group.

[11] Emmanuelle Porcher and Russell Lande. LOSS OF GAMETOPHYTIC SELF-INCOMPATIBILITY WITH EVOLUTION OF INBREEDING DEPRESSION. Evolution, 59(1):46–60, 2005. ISSN 0014-3820. doi: 10.1111/j.0014-3820.2005.tb00893.x. URL https://doi.org/10.1111/j.0014-3820.2005.tb00893.x.

[12] Céline Van de Paer, Pierre Saumitou-Laprade, Philippe Vernet, and Sylvain Billiard. The joint evolution and maintenance of self-incompatibility with gynodioecy or androdioecy. Journal of Theoretical Biology, 371:90–101, 2015. ISSN 0022-5193. doi:10.1016/j.jtbi.2015.02.003. URL https://www.sciencedirect.com/science/article/pii/S0022519315000624.

[13] James M. Bullock, Laura Mallada González, Riin Tamme, Lars Götzenberger, Steven M. White, Meelis Pärtel, and Danny A. P. Hooftman. A synthesis of empirical plant dispersal kernels. Journal of Ecology, 105 (1):6–19, 2017. ISSN 1365-2745. doi:10.1111/1365-2745.12666. URL https://onlinelibrary.wiley.com/doi/abs/10.1111/1365-2745.12666.

[14] E. Porcher and R. Lande. The evolution of self-fertilization and inbreeding depression under pollen discounting and pollen limitation. Journal of Evolutionary Biology, 18(3):497–508, 2005. ISSN 1420-9101. doi:10.1111/j.1420-9101.2005.00905.x. URL https://onlinelibrary.wiley.com/doi/abs/10.1111/j.1420-9101.2005.00905.x.

[15] M. Bartoš, Š. Janeček, P. Janečková, E. Padyšáková, R. Tropek, L. Götzenberger, Y. Klomberg, and J. Jersáková. Self-compatibility and autonomous selfing of plants in meadow communities. Plant Biology, 22(1):120–128, 2020. ISSN 1438-8677. doi: 10.1111/plb.13049. URL https://onlinelibrary.wiley.com/doi/abs/10.1111/plb.13049.

[16] Sarah J. Baldwin and Daniel J. Schoen. Inbreeding depression is difficult to purge in self-incompatible populations of leavenworthia alabamica. New Phytologist, 224(3):1330–1338, 2019. ISSN 1469-8137. doi: 10.1111/nph.15963. URL https://onlinelibrary.wiley.com/doi/abs/10.1111/nph.15963.

[17] Donald M. Waller. Addressing darwin’s dilemma: Can pseudo-overdominance explain persistent inbreeding depression and load? Evolution, 75(4):779–793. ISSN 0014-3820. doi:10.1111/evo.14189. URL https://doi.org/10.1111/evo.14189.

[18] Diala Abu-Awad and Donald Waller. Conditions for maintaining and eroding pseudo-overdominance and its contribution to inbreeding depression. Peer Community Journal, 3, 2023. ISSN 2804-3871. doi:10.24072/pcjournal.224. URL https://peercommunityjournal.org/articles/10.24072/pcjournal.224/.

[19] Deborah Charlesworth and John H. Willis. The genetics of inbreeding depression. Nature Reviews Genetics, 10(11):783–796, November 2009. ISSN 1471-0064. doi: 10.1038/nrg2664. URL https://www.nature.com/articles/nrg2664. xNumber: 11 Publisher: Nature Publishing Group.

[20] Camille Gervais, Diala Abu Awad, Denis Roze, Vincent Castric, and Sylvain Billiard. Genetic architecture of inbreeding depression and the maintenance of gametophytic self-incompatibility. Evolution, 68(11):3317–3324, 2014. ISSN 0014-3820. doi:10.1111/evo.12495. URL https://doi.org/10.1111/evo.12495.

[21] Satoki Sakai. Why are deleterious mutations maintained in selfing populations? An analysis of the effects of early- and late-acting mutations by a two-locus two-allele model. Journal of Theoretical Biology, 533:110956, 2022. ISSN 0022-5193. doi:10.1016/j.jtbi.2021.110956. URL https://www.sciencedirect.com/science/article/pii/S0022519321003751.

[22] Satoki Sakai. Maintenance of high inbreeding depression in selfing populations: Two-stage effect of early- and late-acting mutations. Journal of Theoretical Biology, 502:110307, October 2020. ISSN 0022-5193. doi:10.1016/j.jtbi.2020.110307. URL https://www.sciencedirect.com/science/article/pii/S0022519320301624.

[23] Marcus Frean and Edward R. Abraham. Rock–scissors–paper and the survival of the weakest. Proceedings of the Royal Society of London. Series B: Biological Sciences, 268(1474):1323–1327, 2001. doi:10.1098/rspb.2001.1670. Publisher: Royal Society.

[24] Carol Goodwillie, Susan Kalisz, and Christopher G. Eckert. The Evolutionary Enigma of Mixed Mating Systems in Plants: Occurrence, Theoretical Explanations, and Empirical Evidence. Annual Review of Ecology, Evolution, and Systematics, 36(1):47–79, November 2005. ISSN 1543-592X. doi:10.1146/annurev.ecolsys.36.091704.175539. URL https://www.annualreviews.org/doi/10.1146/annurev.ecolsys.36.091704.175539. Publisher: Annual Reviews.

[25] Michael R. Whitehead, Robert Lanfear, Randall J. Mitchell, and Jeffrey D. Karron. Plant mating systems often vary widely among populations. Frontiers in Ecology and Evolution, 6, 2018. ISSN 2296-701X. doi:10.3389/fevo.2018.00038. URL https://www.frontiersin.org/articles/10.3389/fevo.2018.00038/full. Publisher: Frontiers.

[26] Carly J. Prior and Jeremiah W. Busch. Selfing rate variation within species is unrelated to life-history traits or geographic range position. American Journal of Botany, 108(11):2294–2308, 2021. ISSN 1537-2197. doi: 10.1002/ajb2.1766.

[27] Francisco Encinas-Viso, Andrew G. Young, and John R. Pannell. The loss of self-incompatibility in a range expansion. Journal of Evolutionary Biology, 33(9):1235–1244, 2020. ISSN 1420-9101. doi:10.1111/jeb.13665.

